# Compilation of all known protein changes in the human Alzheimer’s disease brain

**DOI:** 10.1101/2023.04.13.536828

**Authors:** Manor Askenazi, Tomas Kavanagh, Geoffrey Pires, Beatrix Ueberheide, Thomas Wisniewski, Eleanor Drummond

## Abstract

Proteomic studies of human Alzheimer’s disease brain tissue have exceptional potential to identify protein changes that drive disease and to identify new drug targets. Here, we detail a combined analysis of 38 published Alzheimer’s disease proteomic studies, generating a comprehensive map of protein changes in human brain tissue across thirteen brain regions, three disease stages (preclinical Alzheimer’s disease, mild cognitive impairment, advanced Alzheimer’s disease), and proteins enriched in amyloid plaques, neurofibrillary tangles, and cerebral amyloid angiopathy. Our dataset is compiled into a user-friendly, searchable database called NeuroPro. Our combined analysis included 18,119 reported protein differences in human Alzheimer’s disease brain tissue, which mapped to 5,311 total altered proteins. Proteomic studies were remarkably consistent. 848 proteins were consistently altered in ≥5 studies, many of which are understudied in the Alzheimer’s field. Comparison of protein changes in early-stage and advanced Alzheimer’s disease revealed significant synapse, vesicle, and lysosomal changes early in disease, but widespread mitochondrial changes only in advanced Alzheimer’s disease. Comparison of vulnerable and resistant brain regions suggested that protein changes in resistant regions in advanced Alzheimer’s disease are similar to those in vulnerable regions in early-stage Alzheimer’s disease, indicating a temporal progression of protein dysfunction during Alzheimer’s disease advancement. We conclude that NeuroPro is a powerful new resource that provides new insights into human Alzheimer’s disease brain protein changes and highlights novel proteins of particular interest that may mechanistically drive Alzheimer’s disease.

## Introduction

The cause of sporadic Alzheimer’s disease (AD) is currently unknown. Mass spectrometry-based proteomic studies of human brain tissue are an excellent way to uncover the disease mechanisms involved in AD. Protein changes are particularly important to study for a disease such as AD because post-translational events such as protein accumulation, aggregation or post-translational modifications of proteins directly mediate disease ^1^. This has been particularly highlighted by recent studies reporting a poor correlation between mRNA and protein changes in human AD tissue ^2-4^, thus emphasising the need for proteomic studies when examining AD pathogenesis and identifying new biomarkers and/or drug targets.

In recent years there has been a large increase in the number of human AD proteomic studies ^5-7^. These studies have examined protein differences in AD in a variety of brain regions^2-4, 8-29^ and tissue fractions (e.g. insoluble, synaptic, membrane, or blood vessel-enriched fractions^30-35^). Additional studies have performed localized proteomics of neuropathological lesions such as amyloid plaques, neurofibrillary tangles (NFTs), and cerebral amyloid angiopathy (CAA) ^36-42^. Individually, each of these studies have generated important new insight into AD pathogenesis and have uncovered new potential drug targets and biomarkers for AD. Despite these benefits, each of these studies has been somewhat limited when analysed in isolation by either low sample size, inclusion of a limited number of brain regions, or analysis of only one clinical stage of AD.

We hypothesised that a combined analysis of AD human brain tissue proteomic studies would: (1) identify the highest confidence protein changes in AD; (2) resolve potential concerns about inter-study consistency; (3) provide a more comprehensive analysis of AD associated protein changes that could be used to answer key outstanding questions about AD pathogenesis. Therefore, the aim of this study was to perform a combined analysis of mass-spectrometry based proteomic studies examining human AD brain tissue (inclusive of any brain region, any time point in AD, any tissue fraction) to identify consistent protein changes in AD. Inclusion was restricted to studies of human brain tissue, given concerns that animal or cell models do not reflect the complexity of human disease ^26, 40, 43, 44^. We have compiled these data into an online database – NeuroPro – which is a user-friendly resource for the scientific community that details protein changes in three clinical stages of AD (preclinical AD, mild cognitive impairment, and advanced AD), 13 brain regions, and proteins enriched in the three neuropathological hallmarks of AD (amyloid plaques, neurofibrillary tangles, and cerebral amyloid angiopathy). Additionally, we demonstrate the utility of NeuroPro, by using this resource to answer key questions about AD pathogenesis including identification of the earliest protein changes in AD, protein changes associated with selective vulnerability in AD, and correlation of protein enrichment in neuropathological hallmarks and surrounding tissue.

## Results

### Studies included in NeuroPro

38 publications met the inclusion criteria for NeuroPro (**Table 1; Figure 1**). This included 32 studies that identified protein differences in bulk tissue homogenate between AD and controls, 4 studies that identified the proteome of amyloid plaques, 2 studies that identified the proteome of NFTs and 1 study that identified the proteome of CAA. Combined, these studies resulted in 59 unique comparisons of AD vs controls in bulk tissue. The number of individual comparisons was higher than the number of included publications due to some studies examining either multiple brain regions or multiple stages of AD, which each counted as a unique comparison in our analysis. Together, these bulk tissue studies enabled the comparison of protein changes in 13 brain regions and protein changes at three clinical stages of AD (PCL, MCI, AD). A high variance in the number of differentially expressed proteins (DEPs) identified in each study was observed, largely reflecting the sample size in each study. In sum, NeuroPro currently contains data for 18,119 reported protein changes in AD human brain tissue, corresponding to 5,311 significantly altered proteins in AD (**Supplementary Table 1**).

**Table 1:** Studies included in NeuroPro. AD stages refer to cognitively normal controls (“controls”); preclinical AD (“PCL”), mild cognitive impairment (“MCI”) and advanced AD (“AD”). Differentially Expressed Proteins (“DEPs”) shows the total number of all DEPs included in NeuroPro from each study (sum of DEPs from all AD stages and all brain regions). Statistical approach shows the stringency criteria that was used to define DEPs for each study.

| Reference | Brain Region(s) | AD Stage(s) | DEPs included in NeuroPro | Statistical approach |
| --- | --- | --- | --- | --- |
| **Bulk Tissue Homogenate** |  |  |  |  |
| Andreev *et al.* (2012)<sup>8</sup> | Temporal Cortex | Control, AD | 188 | FDR < 5% |
| Astillero-Lopez *et al.* (2022)<sup>16</sup> | Entorhinal Cortex | Control, AD | 399 | FC>1.5 and p<0.05 |
| Bai *et al.* (2020)<sup>26</sup> | Frontal Cortex | Control, PCL, MCI, AD | 20 | FC>1.5 and p<0.05 |
| Carlyle *et al.* (2021)<sup>33</sup> | Parietal Cortex | Control, PCL, AD | 167 | FDR < 5% |
| Dai *et al.* (2018)<sup>17</sup> | Frontal Cortex | Control, AD | 100 | FC>1.5 and p<0.05 |
| Donovan *et al.* (2012)<sup>30</sup> | Frontal Cortex | Control, AD | 9 | FC>1.5 and p<0.05 |
| Hales *et al.* (2016)<sup>31</sup> | Frontal Cortex | Control, PCL, MCI, AD | 308 | FC>1.5 and p<0.05 |
| Haytural *et al.* (2020)<sup>25</sup> | Hippocampus | Control, AD | 65 | FDR < 5% |
| Higginbotham *et al.* (2020)<sup>4</sup> | Frontal Cortex | Control, AD | 102 | FC>1.5 and p<0.05 |
| Ho Kim *et al.* (2015)<sup>12</sup> | Hippocampus | Control, AD | 103 | FDR < 5% |
| Hondius *et al.* (2016)<sup>11</sup> | Hippocampus | Control, AD | 264 | FDR < 5% |
| Johnson *et al.* (2018)<sup>15</sup> | Frontal Cortex | Control, PCL, AD | 841 | AVOVA and Tukey's Post Hoc p<0.05 |
| Johnson *et al.* (2020)<sup>14</sup> | Frontal Cortex | Control, PCL, AD | 1081 | AVOVA and Tukey's Post Hoc p<0.05 |
| Johnson *et al.* (2022)<sup>2</sup> | Frontal Cortex | Control, PCL, AD | 3493 | AVOVA and Holm Post Hoc p<0.05 |
| Li *et al.* (2021)<sup>21</sup> | Occipital Cortex | Control, PCL, AD | 30 | Kruskal-Wallis H test p<0.05 |
| Manavalan *et al.* (2013)<sup>10</sup> | Hippocampus, Cerebellum | Control, AD | 12 | FC>1.5 and p<0.05 |
| McKetney *et al.* (2019)<sup>19</sup> | Entorhinal Cortex | Control, AD | 130 | FC>1.5 and p<0.05 |
| Mendonca *et al.* (2019)<sup>28</sup> | Entorhinal Cortex, Parahippocampal Cortex, Frontal Cortex, Temporal Cortex | Control, MCI, AD | 938 | FC>1.5 and p<0.05 |
| Muraoka *et al.* (2020)<sup>32</sup> | Frontal Cortex | Control, AD | 95 | FC>1.5 and p<0.05 |
| Musunuri *et al.* (2014)<sup>9</sup> | Temporal Cortex | Control, AD | 28 | FC>1.5 and p<0.05 |
| Ojo *et al.* (2021a)<sup>35</sup> | Frontal cortex | Control, AD | 33 | P<0.05 and FC>1.5 AD vs control |
| Ojo *et al.* (2021b)<sup>34</sup> | Frontal cortex | Control, AD | 118 | P<0.05 and FC>1.5 AD vs control |
| Pearson *et al.* (2020)<sup>18</sup> | Ventricle | Control, AD | 25 | FDR < 5% |
| Ping *et al.* (2020)<sup>23</sup> | Frontal Cortex | Control, PCL, AD | 103 | FC>1.5 and p<0.05 |
| Seyfried *et al.* (2017)<sup>3</sup> | Frontal Cortex, Precuneus | Control, PCL, AD | 937 | AVOVA and Tukey's Post Hoc p<0.05 |
| Stepler *et al.* (2020)<sup>24</sup> | Hippocampus, Parietal Cortex | Control, AD | 81 | FC>1.5 and p<0.05 |
| Sweet *et al.* (2016)<sup>13</sup> | Entorhinal Cortex | Control, AD | 103 | FDR < 5% |
| Wang *et al.* (2020b)<sup>46</sup> | Frontal Cortex | Control, AD | 150 | FDR < 5% |
| Wingo *et al.* (2020)<sup>20</sup> | Frontal Cortex | Control, AD | 856 | FDR < 5% |
| Xu *et al.* (2019)<sup>27</sup> | Hippocampus, Entorhinal Cortex, Cingulate Gyrus, Motor Cortex, Sensory Cortex, Cerebellum | Control, AD | 2058 | FDR < 5% |
| Zellner *et al.* (2022)<sup>42</sup> | Parietal cortex | Control, AD | 379 | P<0.05 and FC>1.5 AD vs control |
| Zhang *et al.* (2018)<sup>29</sup> | Frontal Cortex | Control, AD | 269 | FDR < 5% |
| **Amyloid Plaques** |  |  |  |  |
| Drummond *et al.* (2017)<sup>38</sup> | Hippocampus | AD | 1459 | In plaques in >3 cases |
| Drummond *et al.* (2022)<sup>36</sup> | Hippocampus | AD | 2114 | FC>1.5 in plaques vs non-plaques |
| Liao *et al.* (2004)<sup>39</sup> | Frontal Cortex | AD | 22 | FC>1.5 in plaques vs non-plaques |
| Xiong *et al.* (2019)<sup>40</sup> | Hippocampus | AD | 139 | FC>1.5 in plaques vs non-plaques |
| **Neurofibrillary Tangles** |  |  |  |  |
| Drummond *et al.* (2020)<sup>37</sup> | Hippocampus | AD | 542 | In NFTs in >3 cases |
| Hondius *et al.* (2021)<sup>41</sup> | Hippocampus | AD | 112 | FC > 1.5 in NFTs vs control neurons |
| **Cerebral Amyloid Angiopathy** |  |  |  |  |
| Zellner *et al.* (2022)<sup>42</sup> | Parietal cortex | Control, AD | 246 | P<0.05 and FC>1.5 AD vs control |

**Figure 1:**
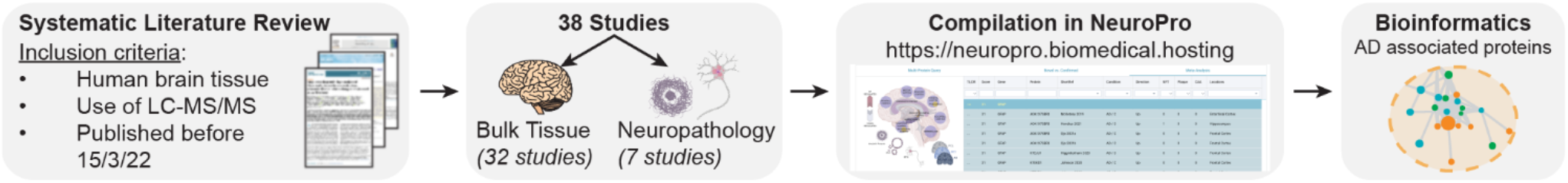
Schematic of methods used in this study.

### Most consistently identified protein changes in AD

54 proteins were identified as DEPs in at least 15 different studies (NeuroPro Score ≥15; **Figure 2B**; **Supplementary Table 1**). The consistent identification of these proteins as significantly altered in AD across so many studies suggests that these protein changes are the most prevalent in AD human brain tissue. 94% (51/54) of these proteins were consistently altered in the same direction across all comparisons: 29 were consistently increased in AD and 22 were consistently decreased in AD. Only 3 proteins were inconsistently altered: VCAN, UCHL1, IDH2. VCAN was predominantly increased in AD bulk tissue studies (11 comparisons) but was decreased in the insoluble brain fraction in MCI and PCL ^31^ and in the CA4/dentate gyrus region of the hippocampus in AD ^12^. IDH2 was predominantly decreased in AD (10 comparisons), but was increased in the insoluble fraction of the frontal cortex in PCL and AD ^31^ and in the wall of the lateral ventricle in AD ^18^. UCHL1 showed a more binary split between studies: it was increased in 5 studies and decreased in 8 studies and there was no obvious reason for this inconsistency; it did not appear linked to differing expression between vulnerable and resistant brain regions, differences between tissue fractions or AD clinical stage.

**Figure 2:**
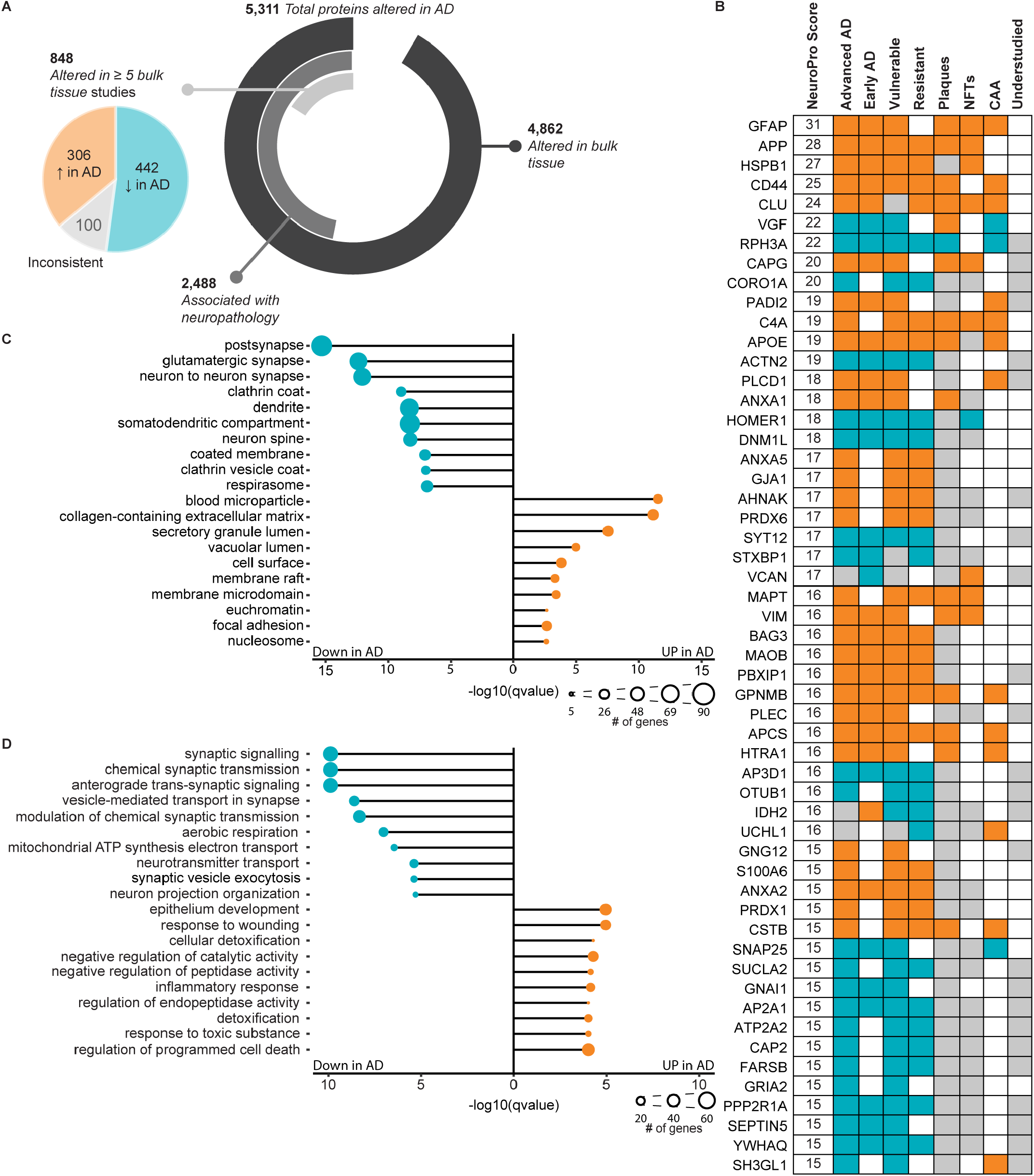
Most consistent protein differences in AD human brain tissue. A: Breakdown of the number of protein differences in NeuroPro. Pie chart shows breakdown of 848 proteins altered in ≥5 bulk tissue studies into consistently increased, consistently decreased and inconsistent subgroups. B: 54 most common protein changes in AD. NeuroPro score: number of studies a protein was significantly altered in. Orange: consistently increased. Blue: consistently decreased. Grey: inconsistently altered. Grey (understudied column): ≤10 studies linking protein with AD. C and D: Most enriched GO terms for proteins (C: cellular component; D: biological process) that are consistently up-regulated (306 proteins) or consistently down-regulated (442 proteins) regulated in AD altered in ≥5 bulk tissue studies.

The most consistently increased proteins in human AD brain tissue were GFAP, APP, HSPB1, CD44 and CLU. The most consistently decreased proteins in human AD brain tissue were VGF, RPH3A, CORO1A, ACTN2 and HOMER1 (**Figure 2B**). These proteins were consistently altered in the same direction across multiple brain regions and often also in preclinical AD and MCI. While some of these most prevalent protein changes were also enriched in neuropathological lesions (**Figure 2B**), this was not always the case, showing that protein enrichment in AD bulk tissue does not necessarily equal enrichment in neuropathological lesions. In addition, there were two instances of proteins that were decreased in AD brain tissue but enriched in neuropathological lesions: VGF and SH3GL1 were both consistently decreased in AD brain tissue, but enriched in plaques and CAA respectively (**Figure 2B**). These results suggest that VGF and SH3GL1 may have unique roles in AD pathogenesis.

Surprisingly, despite consistent detection as DEPs in many AD proteomics studies, 46% of these top 54 proteins are currently understudied in the AD field (classified as ≤10 previous publications linking a protein to AD; **Figure 2B**; **Supplementary Table 1**). In fact, 4 of these top 54 proteins are novel to AD field, with no literature directly linking these proteins to AD; two of these novel proteins were consistently increased in AD (CAPG and PBXIP1) and two were consistently decreased in AD (AP3D1 and SUCLA2). Intriguingly, CAPG is also enriched in both plaques and NFTs, suggesting a potentially important role in pathology. The consistent detection of these proteins as significantly altered in AD proteomic studies warrants future studies examining their role in AD and highlights the power of our combined analysis approach.

### Consistency of Bulk Tissue Proteomic Studies

There was remarkable consistency between bulk tissue proteomic studies, particularly given different sample preparation and mass spectrometry methods, different brain regions, different solubility states and fractionation steps, and different clinical disease stages. 848 proteins were identified as DEPs in AD in ≥5 comparisons of bulk tissue (**Figure 2A**; **Supplementary Table 2**). Of these proteins, there was very high consistency between studies for the direction of change: 306 proteins were consistently increased in AD vs controls and 442 proteins were consistently decreased in AD vs controls. Only 100 proteins showed an inconsistent directional change in AD between studies. It is important to note that inconsistency between studies does not necessarily suggest technical issues, but rather could reflect important pathological protein differences between different stages of disease, different brain regions or in different tissue fractions.

This subset of 848 proteins altered in ≥5 comparisons were highly interconnected (**Figure 3**; p<1.0×10^−16^; protein-protein interaction enrichment). Within this broader protein network, there was evidence of significant enrichment of proteins associated with particular cellular components or biological processes, many of which are known to be associated with AD. For example, there was evidence of significantly decreased synaptic proteins, mitochondrial proteins and vesicle proteins, while there was evidence of significantly increased extracellular and inflammatory proteins (**Figures 2C, D, Supplementary Table 8)**. Our analysis suggested that there was a particularly widespread decrease in many structural components and processes associated with synaptic function including both pre- and post-synaptic proteins, proteins associated with neurotransmitter release (particularly glutamate) and vesicle trafficking (particularly clathrin-coated vesicles). Unexpectedly, our combined analysis also uncovered evidence of decreased kinases associated with tau phosphorylation (including GSK3A, GSK3B, CDK5, MARK1, ROCK2), despite the increased phosphorylation of tau in AD (**Supplementary Table 8)**. There was also strong evidence of increased blood microparticle proteins in AD, and increased proteins associated with neurofibrillary tangles and amyloid plaques.

**Figure 3:**
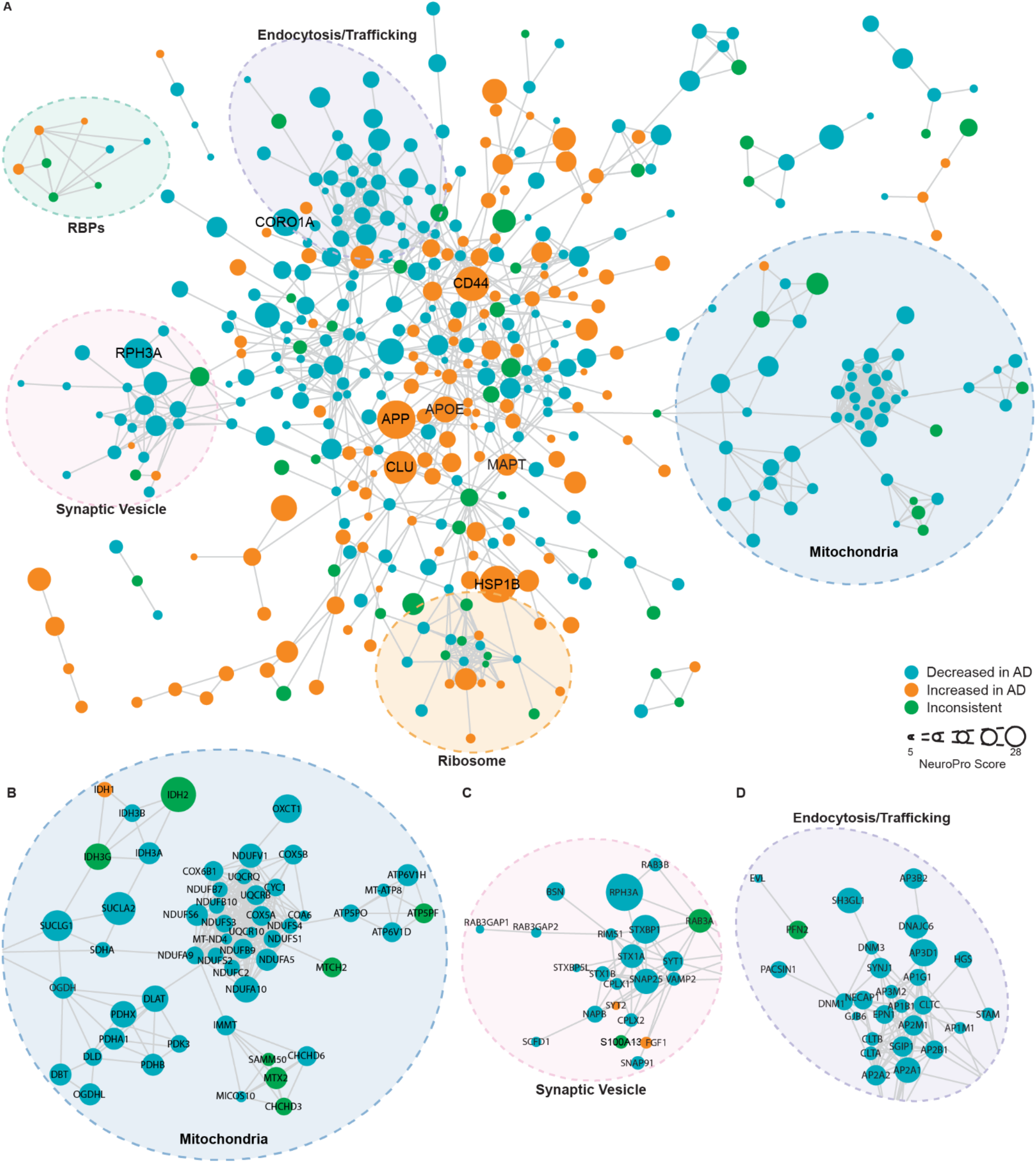
Protein-protein interaction network for 848 proteins altered in ≥5 bulk tissue studies. For simplicity, network shows high confidence (score >0.7), physical interactions only, and only clusters ≥3 proteins shown. Node size reflects NeuroPro score and colour reflects consistent directional change in AD (orange: consistently increased; blue: consistently decreased; green: inconsistently altered). Labelled proteins have a NeuroPro score ≥20 or key AD-associated proteins (APOE, MAPT). Higher magnification of 3 selected sub-networks shown in B-D.

Interestingly, this high inter-study consistency suggests that protein changes in AD consistently occur in the same direction, regardless of brain region or tissue fraction examined. The biggest determinant of inconsistency between studies appeared to be the power of individual studies: as expected, lower powered studies identified only the most extreme protein differences and missed more subtle protein changes. Importantly however, lower power studies did not appear to have a higher prevalence of false positive results, only false negative results, meaning that they were still valuable to include in this combined analysis. The high consistency between studies provided us with increased confidence that NeuroPro could be used to examine fundamental unanswered questions about Alzheimer’s disease pathogenesis. Key examples of how NeuroPro can be used to examine AD pathogenesis are included in the sections below.

### Neuropathology associated proteins

A unique aspect of NeuroPro is that it includes proteomic data examining the three neuropathological hallmarks of AD: amyloid plaques, NFTs and CAA. This allows direct comparison of proteins enriched in each neuropathological lesion for the first time and provides insight into how protein enrichment in neuropathology and more widespread alteration in bulk tissue are related. Studies included in NeuroPro to date identify 2324 plaque proteins, 615 NFT proteins, and 246 CAA proteins (**Supplementary Table 1**). Proteins were classified as enriched or depleted in a neuropathological lesion based on a >1.5 fold change difference when compared to surrounding control tissue. Alternatively, proteins were classified as “present” if they were present in neuropathological lesions in at least 3 cases within a study.

We were most interested to compare proteins that were enriched/depleted in neuropathological lesions as these proteins are more likely to be pathologically relevant. We identified 300 proteins enriched in plaques, 125 proteins depleted in plaques, 54 proteins enriched in NFTs, 58 proteins depleted in NFTs, 192 proteins increased in CAA containing blood vessels and 54 proteins depleted in CAA containing blood vessels (**Figure 4**; **Supplementary Table 1**). Only a small subgroup of proteins were commonly enriched in multiple neuropathological lesions (**Figure 4A**), suggesting that protein enrichment in neuropathological lesions is selective and not simply a result of the “stickiness” of Aβ and tau. Only three proteins were commonly enriched in all neuropathological lesions: C4A, CLU, GFAP. As expected, plaques and CAA showed the greatest number of commonly enriched proteins (37 proteins), including many known amyloid interacting proteins such as APOE, CLU, C3, C4A. One notable exception was APP: APP was not reported as enriched in CAA in the one CAA-specific study included in NeuroPro, contrasting with the large body of evidence confirming that Aβ is the primary component of CAA. This highlights the important caveat that work in this area is still advancing and proteomic results can be significantly influenced by technical factors (e.g. solubilization failure in sample preparation or search parameters). 11 proteins were commonly enriched in plaques and NFTs, likely reflecting the abundant proteins present in phosphorylated tau-rich dystrophic neurites present in neuritic plaques. GO enrichment analysis showed that amyloid plaques were significantly enriched in extracellular matrix and lysosomal proteins (**Figure 4C; Supplementary Table 9**), while NFTs were particularly enriched in neuronal and lumen proteins (**Figure 4D; Supplementary Table 9**). There was almost a complete separation of proteins depleted in neuropathological lesions: only one protein – TARDBP (or TDP43) - was commonly depleted in plaques and NFTs (**Figure 4B**). The fact that all other depleted proteins were unique to each neuropathological lesion suggests that depleted proteins in neuropathological lesions are likely cell environment specific responses to each unique type of neuropathology.

**Figure 4:**
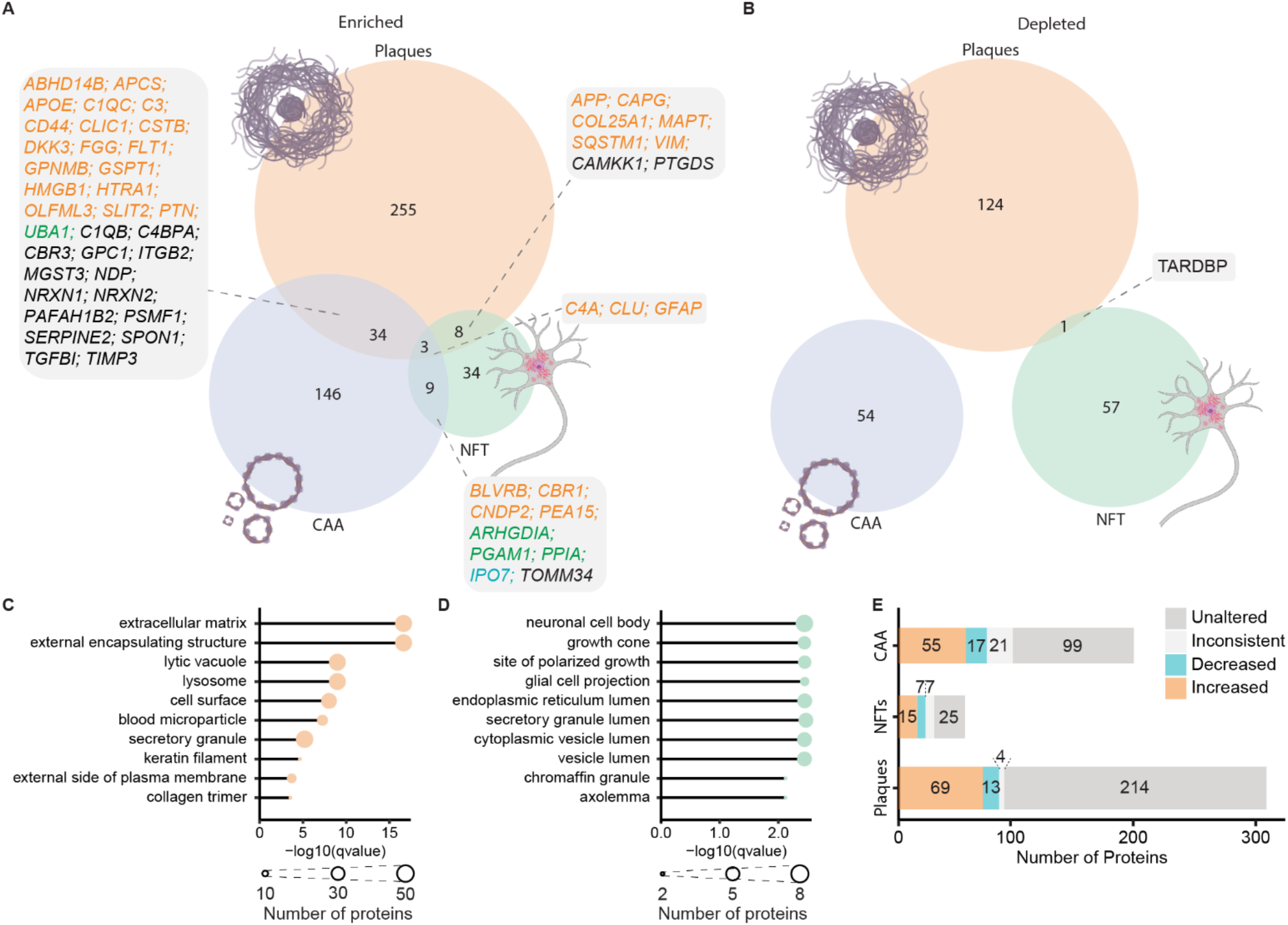
Proteins present in neuropathological lesions. A-B: comparison of proteins enriched and depleted in neuropathological lesions in AD respectively. Boxes specify proteins enriched in multiple lesions; text colour indicates whether proteins are consistently increased (orange), consistently decreased (blue), inconsistently altered (green) in ≥5 bulk tissue studies. Black text indicates proteins either unaltered or altered in <5 studies of AD bulk tissue. C-D: Top enriched GO terms (cellular component) for enriched proteins in amyloid plaques (C) and neurofibrillary tangles (D). Node size shows number of lesion-enriched proteins associated with a GO term. E: Breakdown of directional changes of lesion-enriched proteins in AD bulk tissue. Proteins were increased/decreased if they were consistently altered in ≥5 bulk tissue studies. Proteins were inconsistent if they were inconsistently altered in ≥5 bulk tissue studies. Proteins were unaltered if they were altered in <5 studies of AD bulk tissue.

Comparison of neuropathology enriched proteins and bulk tissue studies showed that neuropathology enriched proteins are not simply those that are also highly enriched in bulk tissue. 214/300 (71%) plaque enriched proteins, 25/54 (46%) NFT enriched proteins and 99/192 (52%) CAA enriched proteins were not consistently altered in ≥5 bulk tissue studies (**Figure 4E**). Intriguingly, there were also a small number of proteins that were consistently depleted in AD in ≥5 bulk tissue studies, while being enriched in neuropathological lesions. This shows that bulk tissue studies cannot be directly used to infer neuropathology-specific changes in AD; while there are some consistencies, this is not always the case.

### Protein Changes at Different Clinical Stages of AD

We were next interested to identify high confidence protein changes that occur in early clinical stages of AD. This is because these protein changes are likely to be initiating drivers of disease and are potentially drug targets for early AD. 15 studies of early AD reached our inclusion criteria for NeuroPro. This included 6 studies of MCI ^26, 28, 31^ and 9 studies of PCL ^2, 3, 14, 15, 21, 23, 31, 33^ (**Figure 5A**). The proteomic data examining protein changes in PCL is particularly robust, having been obtained from multiple high-powered studies. In contrast, the proteomic data currently available for MCI is less comprehensive and was obtained from lower powered studies. Based on this limitation, we defined early AD as either PCL or MCI in our analysis below of early-stage AD proteomic changes.

**Figure 5:**
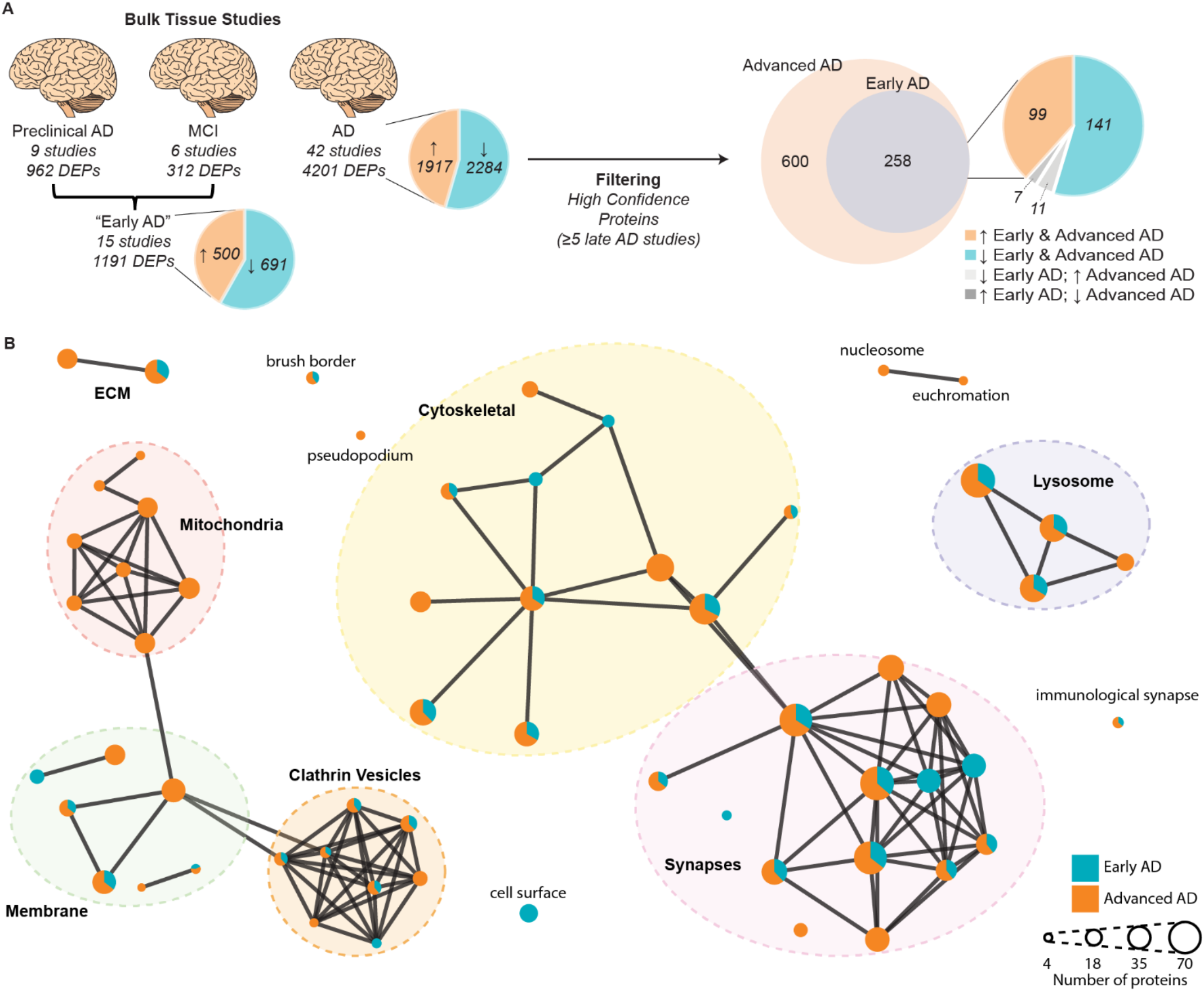
Comparison of protein changes in early-stage and advanced AD. A: Filtering method used to identify high confidence early-stage and advanced AD protein changes. Proteins that were consistently altered in the same direction in each of the three AD clinical stages were identified (DEPs). Inconsistent proteins within a clinical stage were removed. Preclinical AD and MCI were combined to generate an early AD group. Pie charts specify numbers of proteins that are increased/decreased in AD vs controls. Protein changes were filtered to identify high confidence protein changes in early-stage and advanced AD (altered in ≥5 bulk tissue studies). B: GO term (cellular component) network clusters, mapped to show clusters of similar GO terms which were manually annotated to show the overarching cellular component linked to major clusters of GO terms. Each node represents a unique GO term. Node colour shows proportion of proteins for each enriched GO term that were altered in early-stage AD (blue) and advanced AD (orange).

Comparison of the protein changes in early-stage and advanced AD identified 258 protein changes that occurred in both early-stage AD and in ≥5 advanced AD studies. Of these, 240/258 protein changes (93%) occurred in the same direction in both early-stage and advanced AD (**Figure 5A**; **Supplementary Table 3**), suggesting that it is not common for proteins to be increased in early-stage AD and decreased in advanced AD or vice versa. We propose that the 240 protein changes that are consistently altered in the same direction in both early-stage and advanced AD are high confidence early AD protein changes. Of these early-stage AD protein changes, 99 proteins were increased, and 141 proteins were decreased in AD vs controls. This subgroup of early AD proteins was significantly interconnected (p<1.0×10^−16^; Protein-protein interaction enrichment). Pathway analysis particularly highlighted early increases in collagen-containing extracellular matrix proteins (**Supplementary Table 10**). Proteins decreased in early-stage AD were predominantly synapse proteins, which broadly clustered into 3 groups; those associated with the clathrin vesicle coat (most notably strong enrichment of subunits of the AP-2 adapter complex), those associated with synaptic vesicles, and those involved in actin filament organization (**Supplementary Table 10**). Together, this broadly suggests that there is an early synapse dysfunction in AD, predominantly in glutamatergic synapses.

Pathway analysis also showed that most of the earliest affected cellular components (e.g. synapses, cytoskeleton, lysosome and clathrin vesicles) start with changes in a core cluster of proteins in early AD, which then causes a wave of further protein changes in associated proteins as AD progresses (**Figure 5B**). For example, there was strong evidence for exacerbated synapse dysfunction in advanced AD, which started with a core cluster of protein changes in early AD. In particular, there was strong evidence for decreased levels of one type of glutamate receptor - AMPA receptors (evidenced by consistent decreases in 3 of the 4 protein subunits GRIA1, GRIA2, GRIA3) – in advanced AD, but not in early-stage AD. Subunits belonging to other glutamate receptors (Kainate and NMDA receptors) were comparatively less affected in advanced AD (**Supplementary Table 3**). One notable exception to this typical pattern of protein changes through AD progression were widespread changes in mitochondrial proteins, which appeared to be unique to advanced AD (**Figure 5B**). In advanced AD, while decreases in proteins from all 5 complexes of the electron transport chain were observed, this was most prevalent for protein subunits of complex I. Proteins associated with the tricarboxylic acid cycle (TCA cycle), were also selectively decreased in advanced AD, but not early-stage AD (**Supplementary Table 10)**.

Very few proteins were changed in opposite directions in early-stage and advanced AD. 11 proteins were decreased in early-stage AD but increased in advanced AD and 7 proteins were increased in early-stage AD but decreased in advanced AD (**Figure 5A; Supplementary Table 3**). These proteins are particularly interesting as they could represent initially protective protein changes that fail in in later disease stages. Remarkably, 5/7 proteins that were increased in early-stage AD and decreased in advanced AD were mitochondrial proteins (PDHB, DBT, NDUFV1, IDH3G, MMUT), supporting a potential influential mitochondrial role in early-stage AD that is driven by select proteins and not widespread mitochondrial protein changes. The 11 proteins that decreased in early-stage AD but increased in advanced AD showed no enrichment of any pathway, function or cellular component.

### Region specific protein changes in Alzheimer’s disease

Next, we were interested to identify proteins that were linked to selective vulnerability in AD. For this analysis, we used NeuroPro to compare protein changes in advanced AD in 12 brain regions, which were classified as vulnerable or resistant brain regions in AD (**Figure 6A; Supplementary Table 4**). We were particularly interested to determine if there were consistent protein changes in brain regions that are vulnerable and resistant in AD. After filtering for high confidence protein changes, 510 proteins were identified as altered in vulnerable brain regions in advanced AD (**Figure 6A**). 40% (203/510) of these proteins were altered in the same direction in both resistant and vulnerable brain regions (90 proteins were increased in both; 103 proteins decreased in both; **Figure 6B**), providing evidence of AD-associated protein dysfunction in resistant brain regions that do not have widespread neuropathology.

**Figure 6:**
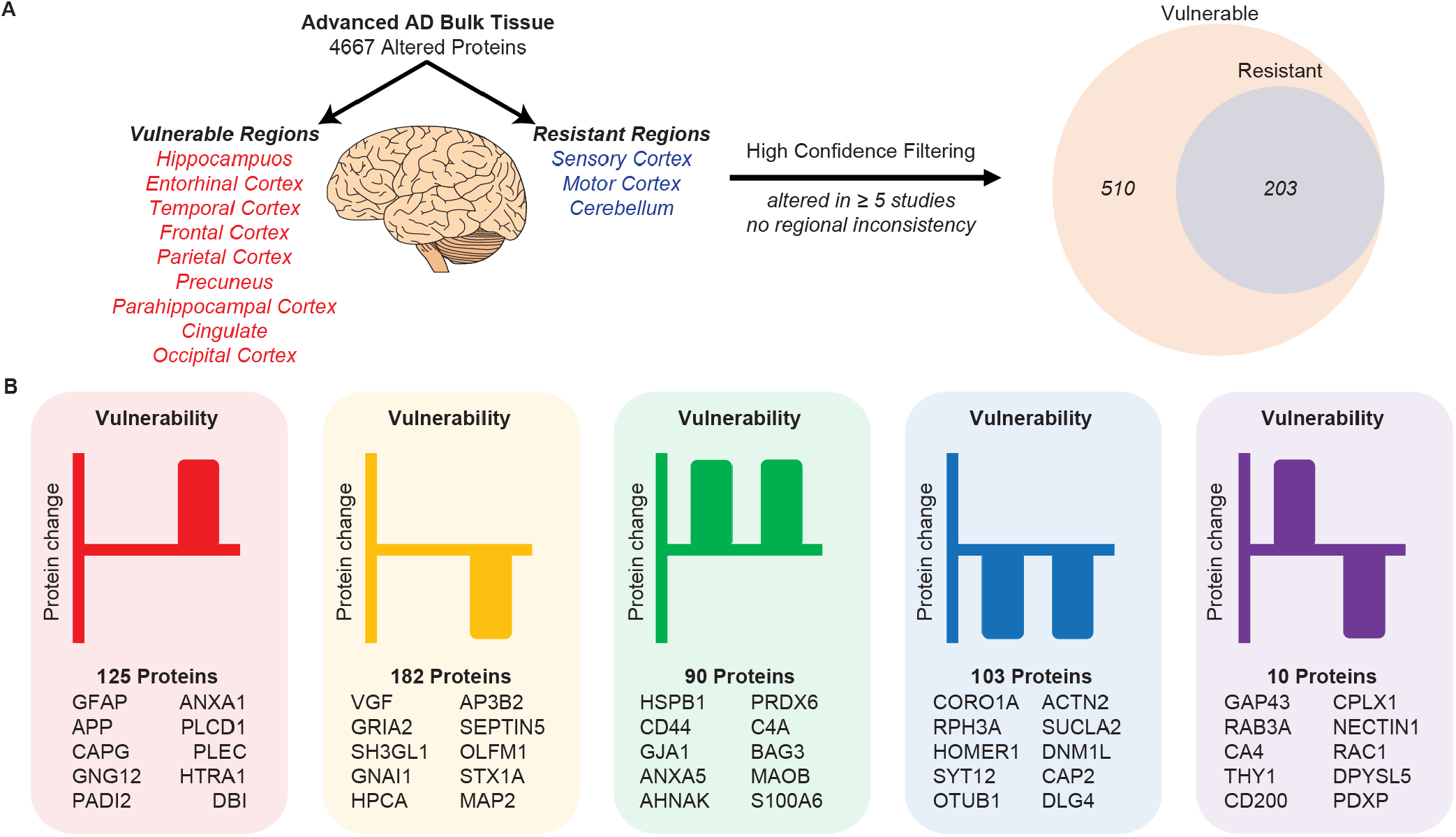
Comparison of protein changes in vulnerable and resistant brain regions. A: Filtering methodology used to identify high confidence protein changes observed in vulnerable and resistant brain regions. Proteins were only considered high confidence protein changes if they were altered in ≥5 bulk tissue studies. B: distribution of protein changes into 5 types of protein changes: proteins increased (red) or decreased (yellow) in vulnerable regions only, proteins increased (green) or decreased (blue) in both vulnerable and resistant regions, proteins increased in resistant regions and decreased in vulnerable regions (purple). Proteins with the highest total NeuroPro score in each group are specified.

21% (64/307) of protein changes that were uniquely present in vulnerable brain regions, but not resistant brain regions were proteins enriched in neuropathological lesions (e.g. GFAP, APP, VGF, HTRA1, MDK, SMOC1, SQSTM1), confirming that neuropathology is not a predominant feature in low vulnerability regions in advanced AD (**Supplementary Table 4**). Very few proteins showed opposite protein changes in vulnerable regions vs resistant regions (10 proteins; all decreased in high/moderate vulnerable regions but increased in low vulnerable regions).

Based on the significant overlap in protein changes in vulnerable and resistant brain regions in advanced AD, we hypothesized that protein changes in resistant brain regions in advanced AD may be the same as those in vulnerable regions in early-stage AD. If so, this would reflect a temporal wave of progressive protein dysfunction through affected brain regions as AD progresses. To test this hypothesis, we directly compared protein changes in resistant regions in advanced AD with protein changes in vulnerable regions in early-stage AD (**Supplementary Table 5**). This analysis showed considerable overlap between these two groups of proteins, supporting our hypothesis. 64 proteins were altered in the same direction in the two datasets (**Figure 7A**; **Supplementary Table 5**); 35 proteins were decreased in AD and 29 proteins were increased in AD. We propose that these protein changes are some of the earliest protein changes in AD, occurring prior to the development of neuropathology and persisting throughout disease progression. Notably, while this subset of “pre-neuropathology protein changes” includes many well-known AD-associated proteins (e.g. APOE, MAOB, AQP4), it does not include APP and MAPT. This reflects the fact that widespread neuropathology is not yet present in resistant brain regions. Pathway analysis of pre-neuropathology protein changes highlighted increased levels of chaperones associated with aggregated Aβ and tau (HSPB1, HSPB8, BAG3, APOE, APCS), increased levels of enzymes associated with energy production and biosynthesis of neurotransmitters (SPR, PKM, PAICS, MTAP, MAOB, LTA4H, GYG1, BBOX1, ALAD) and proteins involved in innate immunity. A significant subgroup of these pre-neuropathology protein changes were synaptic proteins (24/64 proteins; **Figure 7B**). These altered synaptic proteins were associated with many intracellular components in both the pre- and post-synapse (**Figure 7B**), suggesting widespread early synaptic dysfunction in AD. Based on this analysis we propose that there are three phases of protein changes in AD human brain tissue: Phase 1 (pre-neuropathology protein changes), Phase 2 (Early-stage AD protein changes which occur early in disease alongside neuropathology development), and Phase 3 (Advanced AD protein changes) (**Supplementary Table 6**).

**Figure 7:**
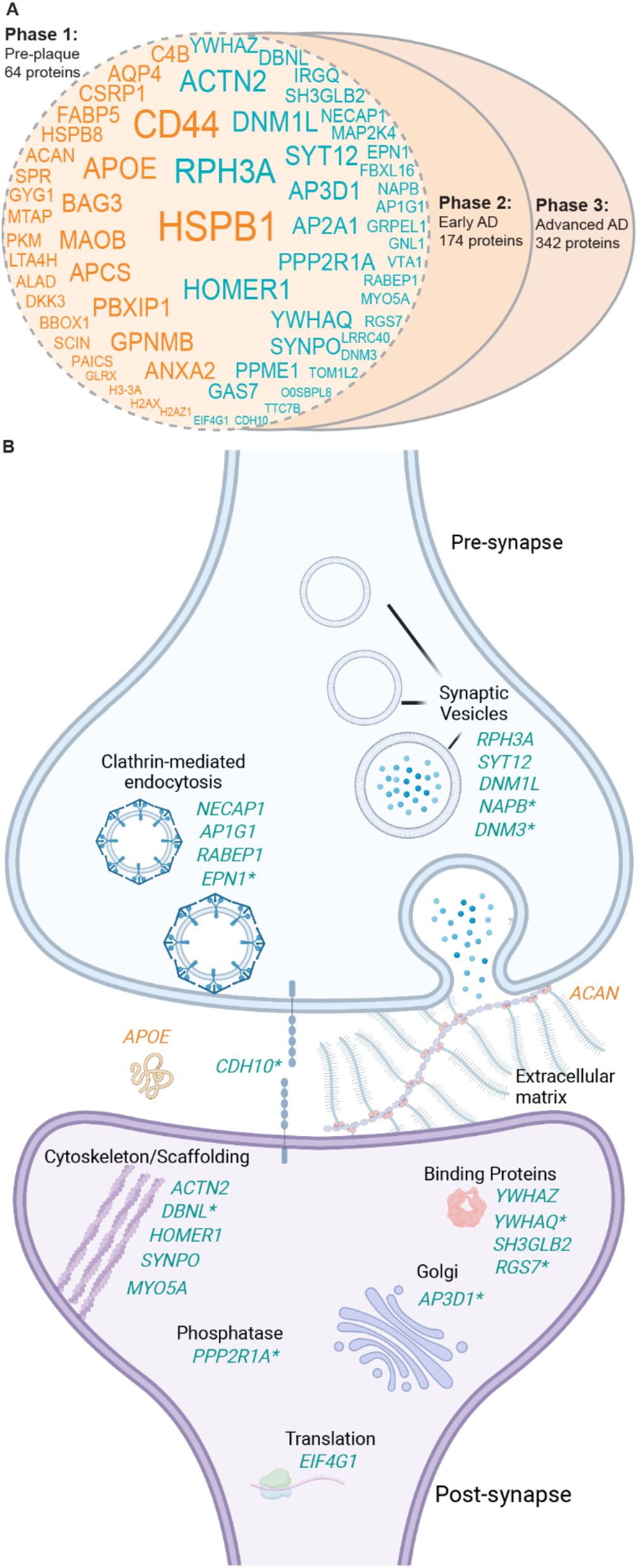
Proposed “pre-neuropathology” protein changes. A: High confidence protein changes were mapped to three successive phases of disease: pre-neuropathology phase, early AD phase, and advanced AD phase. The 64 proposed pre-neuropathology protein changes are identified by gene IDs. Text size reflects total number of studies each protein has been reported to be significantly altered in (i.e. NeuroPro score). Orange text: proteins consistently increased in AD vs controls; blue text: proteins consistently decreased in AD vs controls. B: 24 pre-neuropathology protein changes were synaptic proteins. Schematic highlights simplified key locations of each of these 24 altered proteins based on GO terms, noting the caveat that many of these proteins have multiple functions and locations within the synapse, which is not represented here. 22 synaptic proteins were down-regulated (blue text), while two synaptic proteins were up-regulated (orange text). * indicates proteins that are reported to be located in both the pre- and post-synapse. For simplification, each protein is only highlighted once in either the pre- or post-synapse.

## Discussion

NeuroPro provides a comprehensive roadmap of protein changes that occur in the human brain throughout the progression of AD. NeuroPro provides new insights into AD pathogenesis and highlights new potential drug targets and biomarkers for AD. As such, we propose that it is a useful innovative resource for the AD field. Its novelty lies in our combined analysis approach of diverse mass spectrometry datasets that often have limited power when analysed in isolation. Our online database allows users to immediately place AD brain protein changes in the context of clinical disease stage, brain region specificity, association with neuropathology and subcellular/biochemical changes in disease (e.g. subcellular localization or insolubility in disease). To demonstrate the power of NeuroPro, we have used it here to examine key questions about AD pathogenesis. In doing so, we have: (1) shown that proteomic studies of AD tissue are highly consistent, (2) shown that proteins enriched in plaques, NFTs and CAA are largely unique and independent of broader protein changes in bulk tissue, (3) shown that there are many similar protein changes in resistant brain regions in AD and early clinical stages of AD (4) identified some of the earliest protein changes in AD, including a subset that we hypothesise to occur prior to neuropathology development. These results are just a few examples of the type of analyses that can be performed using NeuroPro. An additional key benefit of NeuroPro is that users can search their own datasets. NeuroPro provides immediate context to newly generated data about the involvement of protein hits in AD and allows rapid comparison between protein changes in AD and other diseases.

One of our key findings was that widespread decreases in synapse proteins appears to be one of the earliest pathological changes observed in human AD brain tissue. While synaptic changes are known to have a pivotal role in AD ^47, 48^ and have been reported in both preclinical AD and MCI ^49-51^, exactly how early synaptic dysfunction occurs in human AD and the initiating protein drivers involved are unknown. We identified a sub-group of 24 synapse proteins that were significantly altered in resistant brain regions prior to the development of local AD neuropathology, suggesting that these synaptic protein changes could be among the first pathological changes in AD. We hypothesize that these initial synaptic protein changes promote downstream synaptic dysfunction in neighbouring proteins, interactors or members of the same networks as AD progresses, which is reflected in the increased number of synaptic protein changes at later stages of AD. These pre-neuropathology synapse protein changes were observed throughout both the pre- and post-synapse and were associated with many subcellular compartments, suggesting widespread dysfunction throughout the synapse in early AD. While little is known about how most of these proteins are mechanistically involved in AD, the role of some of these proteins in AD has been previously explored: for example, DNM1L^52, 53^, SYNPO^54^ and YWHAZ^55, 56^ all appear to promote AD-associated pathology. Mechanistic roles for other non-synaptic protein changes in this “pre-neuropathology” phase have also been reported, strengthening support for the potential importance of this subset of proteins. For example, there is evidence that HSPB1^57^, BAG3^58^, GPNMB^59^, AQP4^60^, HSPB8^61, 62^, PKM^63^, DKK3^64^, GLRX^65^, GAS7^66^ are protective in AD, while MAOB^67^ and CD44^68^ appear to promote AD-associated pathology. These previous studies confirm that many of these pre-neuropathology protein changes are potential drivers of AD and suggest that the mechanistic roles of the remaining proteins in this group should be examined in future studies.

Another key finding was that enrichment of proteins in neuropathological lesions is a selective process that is unique to each specific type of lesion. We hypothesize that the enriched proteins present in each type of lesion reflect the processes involved in lesion development. For example, plaques were significantly enriched in lysosomal proteins, nicely complementing recent studies that suggest that amyloid plaques form after accumulation of intraneuronal Aβ in autophagic vacuoles ^69^. In contrast, NFTs were significantly enriched in neuronal and endoplasmic reticulum proteins, supporting previous studies showing a strong association between tau and ribosomal proteins that can pathologically impair translation ^70, 71^. In addition, our results highlight the importance of localized proteomics approaches^72-74^ to identify neuropathology enriched proteins, as protein enrichment in neuropathology was often not reflected in bulk tissue studies.

We were intrigued to see that many AD-associated protein changes were consistently observed in the same direction in early AD, advanced AD, vulnerable brain regions and resistant brain regions. Together, our results suggest that resistant brain regions in advanced AD may develop the same protein changes observed in vulnerable brain regions in early AD; supporting a hypothesis that there may be a temporal wave of progressive protein changes throughout the brain in AD. While it is well established Aβ and tau progressively spread through the brain in AD ^75, 76^, here we show that this same process may occur for many other proteins also. Importantly, a subset of these protein changes (many of them synaptic proteins) appears to occur before Aβ and tau accumulation, suggesting that they may be early disease drivers. Our results support the concept that resistant brain regions are initially protected against pathology (possibly due to a range of factors ^77-79^); however, once these protective measures eventually fail, the same initiating protein changes observed in vulnerable brain regions in early AD become apparent. If true, this has potential implications for experimental studies as resistant brain regions in advanced AD could be used as a proxy to study early AD-associated protein changes in human brain tissue. Given these implications, these results should be further explored in future studies to confirm our preliminary findings proposed here.

There are several limitations to our study, which highlight critical future research directions. For example, only a limited number of MCI proteomic studies were available, and these were largely underpowered.

Future, high-powered proteomic studies comparing protein changes in MCI and PCL tissue are needed, as these could be used to specifically identify protein changes linked to cognitive impairment while controlling for neuropathology. Furthermore, most AD proteomics studies to date have examined protein changes in frontal cortex: only a handful have examined other brain regions. Additional high-powered studies examining other brain regions, particularly resistant brain regions, would be exceptionally useful to determine the temporal progression of protein changes throughout the progression of AD. In addition, more studies are required to confirm proteins enriched in neuropathological lesions. While proteins that are “present” in neuropathological lesions are interesting, proteins that are selectively enriched in neuropathological lesions in comparison to neighbouring control tissue are more likely pathology drivers. In particular, additional datasets examining CAA are needed given that the current single dataset that met our inclusion criteria did not identify Aβ and other major amyloid associated proteins ^42^, which we propose to be due to the lack of solubilization of Aβ in these samples because of the lack of formic acid treatment during sample preparation prior to mass spectrometry in this study.

To conclude, we have shown that NeuroPro is a powerful new resource that provides novel insights into AD pathogenesis and highlights many novel or understudied proteins in the AD field, providing exciting avenues of research for future studies. This has the potential to increase the impact and widespread use of proteomic data and will hopefully pave the way forward for new therapies and biomarkers for AD.

## Online Methods

### Systematic literature review and inclusion/exclusion criteria for NeuroPro

A comprehensive literature search was performed to identify all LC-MS/MS studies of human AD brain tissue. The following search terms were used on Pubmed: “Alzheimer’s proteomics” and “mass spectrometry human brain Alzheimer’s”. All papers published prior to March 15^th^ 2022 were included in the search. Search results were manually filtered to identify acceptable studies that defined protein changes in bulk tissue homogenate between (i) AD vs controls (ii) mild cognitive impairment (MCI) vs controls (iii) preclinical AD (PCL; also referred to in the literature as asymptomatic AD, prodromal AD or high pathology control) vs controls. Inclusion criteria for bulk tissue homogenate studies were: analysis of human brain tissue, use of LC-MS/MS, and sufficient accessible LC-MS/MS data. Proteins were considered to be differently expressed between disease and control (DEPs) based on any of the following statistical approaches: FDR of <5%, p<0.05 using ANOVA and an appropriate post-hoc test (e.g. Tukey’s or Holm’s comparison post hoc test), p<0.05 using Kruskal-Walis H test, or a combination of p<0.05 using t-test combined with a fold change difference >1.5 fold. Studies using 2D-gel electrophoresis were excluded. Studies that did not specify brain region (e.g. analysed proteins in “cortex”) were excluded. Studies that did not provide a protein identifier (or an adequate identifier that could be used to map a corresponding protein identifier) in their datasets were excluded.

Datasets were then manually adjusted to permit direct comparison between studies using the following methods: Single gene ID and UniProt ID were generated for each reported protein; if multiple gene IDs for a single protein were provided, the first listed Gene ID was used. UniProt IDs were stripped of isoforms.

Duplicate gene IDs within a dataset were removed. p-values (generated using unpaired t-test) and fold change differences between AD and control groups were manually performed using published data if not provided in the original study and sufficient data was available. Proteins identified by only 1 peptide were excluded. Published lists of DEPs were manually filtered to only include those that reached our stringency criteria detailed above.

Proteomics studies examining the proteome of amyloid plaques, neurofibrillary tangles (NFTs) or cerebral amyloid angiopathy (CAA) were also included. Inclusion was limited to studies that selectively isolated neuropathological lesions using either laser capture microdissection (for plaques and NFTs) or a dextran/Ficoll gradient extraction (for CAA containing blood vessels). A protein was considered “present” in plaques or NFTs if it was present in ≥3 cases within a study. Reported results were manually filtered to exclude proteins identified by only 1 peptide. Proteins were considered “increased” or “decreased” in a plaque, NFT, or CAA containing blood vessel based on >1.5 fold change difference between plaques:non-plaque regions, NFT containing neurons:non-NFT containing neurons or CAA containing blood vessels:non-CAA containing blood vessels.

### NeuroPro Database

Data from selected studies were uploaded into NeuroPro: https://neuropro.biomedical.hosting. All proteins were annotated with both a protein identifier (UniProt ID) and gene ID. Proteins were grouped within NeuroPro using GeneID. The entire NeuroPro dataset can be downloaded in the “Meta-Analysis” tab within the NeuroPro database. A “NeuroPro Score” for each protein was generated based on the number of times that protein was reported as significantly altered in AD in published studies. In NeuroPro, proteins can be filtered according to brain region, disease stage, association with neuropathology or direction of change in AD, either alone or in combination in the “meta-analysis” tab. Resulting filtered datasets can be exported. Users can also directly compare datasets of interest (e.g. user-generated data or published proteomic data examining a different disease) in the “Multi-Protein Query” tab, which categorizes searched proteins into known or novel AD-associated proteins in the “Novel vs. Confirmed” tab.

### Analysis of NeuroPro

#### Consistency of directional change

Data was exported from NeuroPro for analysis on 7-21-22 (**Supplementary Table 1**). The NeuroPro score for each protein was obtained from NeuroPro and an additional NeuroPro (Bulk Tissue) Score was generated, which was a count of the number of times a protein was designated a DEP in studies of bulk tissue homogenate only (i.e. excluding studies of neuropathological lesions). Proteins were designated as “Increased/Decreased in AD” if they were consistently altered in the same direction in ≥5 studies of bulk tissue homogenate, with one outlier permitted (to accommodate the range of brain regions, fractions, and disease stages). DEPs in ≥5 studies of bulk tissue homogenate with two or more outlier directional changes were designated “Inconsistent in AD”.

#### Comparison of protein changes in early-stage vs advanced AD

Bulk tissue data were exported from NeuroPro. DEPs were filtered into those occurring in PCL, MCI and advanced AD and individual NeuroPro Scores were assigned for each clinical stage of AD. Proteins were designated as “Increased” or “Decreased” in each clinical stage of AD if they were consistently altered in the same direction in all studies within that clinical stage of AD. One directional outlier was permitted only in cases where a protein was altered in ≥5 studies within a clinical stage. Proteins that were increased in AD received a positive score and proteins that were decreased in AD received a negative score. All other proteins were classified as having “Inconsistent” directional change within that clinical stage of AD and did not receive a score. The resulting dataset was then filtered to only contain proteins with a score in at least one stage of AD (**Supplementary Table 3**).

Protein changes were compared in early-stage AD (defined as either PCL or MCI) and advanced AD. Advanced AD protein changes were restricted to include high confidence protein changes only, which were classified as consistently present in ≥5 studies of advanced AD (one directional outlier permitted). Protein changes were grouped into those that were (1) altered in the same direction in both early-stage and advanced AD (proposed “early-AD protein changes”), (2) uniquely altered in advanced AD and not early-stage AD (proposed “advanced AD protein changes”), (3) altered in the opposite direction in early-stage and advanced AD or altered only in early-stage AD and not in advanced AD (proposed “opposite protein changes in advanced AD and Early-AD”). All other proteins were classified as “Unable to group”. This dataset was then compared to MitoCoP – a dataset of high confidence human mitochondrial proteins ^45^ and SynGo – a dataset of high-confidence human synaptic proteins (2021 download; https://www.syngoportal.org/), to highlight mitochondrial and synaptic proteins respectively.

#### Comparison of protein changes in different brain regions

Bulk tissue data examining advanced AD was exported from NeuroPro. This analysis was restricted to advanced AD as current proteomic studies of early-stage AD tissue have largely been restricted to the frontal cortex, therefore precluding an in-depth regional comparison. Advanced AD proteomic data was available for 13 brain regions; frontal cortex, hippocampus, parahippocampal cortex, entorhinal cortex, temporal cortex, parietal cortex, cingulate gyrus, precuneus, motor cortex, sensory cortex, occipital cortex, ventricle wall and cerebellum. Brain regions were designated as either vulnerable or resistant in AD based on consensus in the literature about the timing and extent of neuropathology. Vulnerable regions included: entorhinal cortex, hippocampus, parahippocampal cortex, temporal cortex, frontal cortex, parietal cortex, precuneus, cingulate gyrus, occipital cortex. Resistant regions included: sensory cortex, motor cortex, cerebellum. Data examining proteomic changes in the ventricle wall^18^ were not included in this analysis as there is not sufficient literature available to determine whether this region is vulnerable or resistant in AD. A brain region specific NeuroPro score was generated for each region, which was a count of the number of studies where that protein was reported to be significantly altered in AD within that brain region. Proteins consistently increased in AD vs controls received a positive score and proteins consistently decreased in AD vs controls received a negative score.

An analysis of protein differences in Vulnerable and Resistant brain regions was performed (**Supplementary Table 4**). High confidence protein differences were identified as those with a combined region score of ≥5. These high confidence protein changes were then categorized as; (1) Increased in Vulnerable; Unchanged in Resistant, (2) Decreased in Vulnerable; Unchanged in Resistant, (3) Increased in Vulnerable and Resistant (4) Decreased in Vulnerable and Resistant, (5) Increased in Vulnerable; Decreased in Resistant, (6) Decreased in Vulnerable; Increased in Resistant, (7) Inconsistent.

#### Analysis of the pre-neuropathology AD protein changes

“Pre-neuropathology protein changes” were designated as those consistently present in same direction in resistant brain regions, early-stage AD and advanced AD (**Supplementary Tables 5 and 6**). This subset of pre-neuropathology protein changes was identified by direct comparison of early-AD protein changes (**Supplementary Table 3**) and protein changes in resistant brain regions (**Supplementary Table 4**).

Proteins that were consistently altered in both early-stage AD and advanced AD, but not in resistant brain regions were designated as “Early AD protein changes” and protein changes that were consistently altered only in advanced AD were designated as “Advanced AD protein changes”. Protein changes that were altered in an inconsistent direction between any of the protein subsets analysed (resistant protein changes; early AD protein changes; advanced AD protein changes) were classified as inconsistent protein changes and excluded from analysis (**Supplementary Table 7)**.

### Pathway/Network Analysis or comparison with previous datasets

General data manipulations and grouping were performed in R v4.0.2 using the tidyverse v1.3.2 collection of packages. Plots were generated in R with the packages ggplot2 v3.4.0, ggpubr v0.5.0, ggrepel v0.9.3, and all figures were edited in Adobe Illustrator v27.1.1. Gene Ontology (GO) enrichment analysis was performed in R using the packages enrichplot v1.16.2, clusterProfiler 4.4.4, using the genome wide annotation for human, org.Hs.eg.db v3.15.0. Prior to analyses gene IDs were mapped to Entrez IDs with the ‘bitr’ function of clusterProfiler. GO terms were filtered to an FDR < 0.05 and the full list of proteins detected were used as the background list (5,311 proteins). Primarily the GO cellular compartment (GOCC) and GO biological process (GOBP) annotations were used. GO terms were reduced with the ‘simplify’ function from clusterProfiler to reduce heavily redundant terms prior to plotting. GO terms were plotted as either top 10 ‘lollypop’ plots or enrichment maps where nodes (GO annotations) are connected by shared proteins. Protein–protein interaction networks were generated in STRING v11.5 and the networks were edited in Cytoscape v3.9.1 and Adobe Illustrator. Pathway collections were annotated manually based on string gene ontology outputs. Venn diagrams were generated with the R package Venerable v 3.1.0.9000 and edited in Adobe Illustrator. Selected figure panels were created with BioRender.com.

### Comparison with previous literature

Systematic Pubmed searches were performed to determine a particular protein’s known association with AD. Search terms used were ““protein name” or “gene ID” Alzheimer’s” and the search was performed on 15-9-22. Protein name was obtained from UniProt.

## Acknowledgements

This study was supported by funding from NIH (P01AG060882 and P30AG066512), Bluesand Foundation, Alzheimer’s Association (Blas Frangione Early Career Award), and the National Health and Medical Research Council of Australia (Program Grant # 1132524).

## Data Availability

All data used in this study is available on the open-access NeuroPro website: https://neuropro.biomedical.hosting. All data used in sub-analyses are included in full in the supplementary tables of this manuscript.

## Author Contributions

E.D. conceived and designed this study and was responsible for identification of studies. E.D and M.A. were responsible for data curation and combined analysis. M.A. designed and maintains the NeuroPro database. T.K. and E.D. were responsible for bioinformatics analyses and figure generation. G.P., T.W., B.U. provided expert advice on the interpretation of data. E.D. wrote the manuscript with input from co-authors. All authors read and approved the final manuscript.

## Supplementary Tables

See readme tab in the excel file.

